# Extracellular matrix gene expression signatures as cell type and cell state identifiers

**DOI:** 10.1101/2021.03.11.434939

**Authors:** Fabio Sacher, Christian Feregrino, Patrick Tschopp, Collin Y. Ewald

## Abstract

Transcriptomic signatures based on cellular mRNA expression profiles can be used to categorize cell types and states. Yet whether different functional groups of genes perform better or worse in this process remains largely unexplored. Here we test the core matrisome - that is, all genes coding for structural proteins of the extracellular matrix - for its ability to delineate distinct cell types in embryonic single-cell RNA-sequencing (scRNA-seq) data. We show that even though expressed core matrisome genes correspond to less than 2% of an entire cellular transcriptome, their RNA expression levels suffice to recapitulate important aspects of cell type-specific clustering. Notably, using scRNA-seq data from the embryonic limb, we demonstrate that core matrisome gene expression outperforms random gene subsets of similar sizes and can match and exceed the predictive power of transcription factors. While transcription factor signatures generally perform better in predicting cell types at early stages of chicken and mouse limb development, *i.e.,* when cells are less differentiated, the information content of the core matrisome signature increases in more differentiated cells. Our findings suggest that each cell type produces its own unique extracellular matrix, or matreotype, which becomes progressively more refined and cell type-specific as embryonic tissues mature.

**Highlights:**

- Cell types produce unique extracellular matrix compositions
- Dynamic extracellular matrix gene expression profiles hold predictive power for cell type and cell state identification

## Introduction

How to define and identify different cell types remains a fundamental challenge in biology [1–4]. Cell types have traditionally been classified based on their morphology and function, by the tissues from where they were isolated, their ontogenetic origin, or their molecular signatures [3]. In recent years, gene expression data from single-cell transcriptomic studies (scRNA-seq) have been used to characterize and fine-tune different cell type classification systems [2,3,5].

Cellular fate and cell-type-specific gene expression programs are thought to be largely regulated by transcription factors and their corresponding *cis*-regulatory networks [2,4,6]. Accordingly, transcription factor expression profiles can be useful in identifying cell types from scRNA-seq data [2,7,8]. Yet other cellular properties can also vary dynamically, in a cell type-specific manner. Hence, we were looking for additional sets of putative ‘biomarker’ genes, to identify cell types and states.

The extracellular matrix (ECM) has traditionally been thought of as a static protein network surrounding cells and tissues. However, the ECM has recently emerged as a highly dynamic system [9–11]. In fact, transcription and translation of some ECM genes are even coupled to circadian rhythm, highlighting the dynamic nature of ECM composition [12]. Experimentally, ECM composition has so far been determined mostly by proteomics assays [13]. More recently, *in-silico* approaches have defined the ‘matrisome’ gene sets representing all genes either forming or remodeling the ECM, as present in a given species’ genome [13,14]. The matrisome is divided into two main categories: the core matrisome encompassing all proteins that form the actual ECM (collagens, glycoproteins, proteoglycans) and the matrisome-associated proteins that either bind to the ECM, remodel the ECM, or are secreted from the ECM [13,14].

Importantly, it has been postulated that each cell type produces its own unique ECM [14– 17]. To capture this concept, we have recently defined the ‘matreotype’, an extracellular matrix signature associated with -or caused by -a given cellular identity or physiological status [17]. For instance, cellular status, including metabolic, healthy or pathologic, or aging have been associated with distinct ECM expression patterns (*i*.*e*., matreotypes) [14,17–21]. Furthermore, cancer-specific cell types can be identified based on their unique ECM composition [13,14,20,22]. This indicates that ECM composition is plastic and adapts to cellular needs or status. Since this is a highly dynamic process, snapshots of unique ECM compositions are reflected in distinct matreotypes.

Based on this, we hypothesized that ECM gene expression is a dynamic parameter that could hold predictive value to function as a biomarker for cell type and state identification. To test our hypothesis, we re-analyzed publicly available scRNA-seq data and specifically examine ECM gene expression signatures. Unsupervised clustering of scRNA-seq data using the whole transcriptome -or highly variable genes therein -is a common strategy to classify cell types [2,3,5]. Here we use defined transcriptome subsets -namely, expressed transcription factors, core matrisome genes, and random transcriptome subsets of equal size -to re-cluster scRNA-seq data and evaluate the resulting clusters in comparison to the performance of the entire transcriptome. In embryonic data coming from chicken and mouse limbs, we find that the core matrisome has less predictive power in undifferentiated cells, early during development, but outcompetes transcription factors later in development and in more differentiated cell types. Consequently, we propose matreotype gene expression signatures as context-dependent proxies for identifying cell types.

## Results

### Defining the chicken core matrisome

The matrisome has been defined for humans (1027 genes), mice (1110 genes), zebrafish (1002 genes), planarian (256 genes), *Drosophila* (641 genes), and *C. elegans* (719 genes), where it corresponds to roughly 4% of their protein-coding genes [14,23–26]. In order to expand the number of model organisms amenable to ‘matreotype’ investigation, we first decided to define the chicken matrisome. Using the 1110 mouse and 1027 human matrisome gene lists to perform orthology and InterPro domain searches, we identified 631 and 656 chicken matrisome genes, respectively (Supplementary Fig. 1, Supplementary Table 1). In summary, we define the chicken matrisome with 217 core-matrisome genes and 443 associated-matrisome genes (Supplementary Table 1).

### The chicken core matrisome as a molecular signature with cell-type specificity

To evaluate the cell type clustering performance of the ‘chicken core matrisome’, we re-analyzed embryonic stage HH29 (stage 29 Hamburger and Hamilton) [27] chicken hind limb scRNA-seq data [28]. At this point of development, chicken limb progenitor cells have already differentiated into transcriptionally distinct tissue types [28], which is reflected in the separation of our t-distributed Stochastic Neighbor Embedding (t-SNE) dimensionality reduction and the superimposed, color-coded clustering information (Fig. 1A). We compared the cell type clustering of the core matrisome to the entire transcriptome and contrasted its performance with highly variably expressed transcription factors -representing a ‘traditional cell type identifier’ -and an equal number of randomly picked genes, to estimate baseline clustering. Of the 217 chicken core matrisome genes, 136 were expressed in our limb scRNA-seq data (Data Source File 1). Accordingly, we picked 136 genes randomly, as well as the 136 most variably expressed transcription factors, chosen by maximum variance across all cells in the sample. With these three small subsets of genes -representing only 1.26% of all expressed genes -, we re-clustered our data using the Louvain-Jaccard algorithm. We adjusted the resolution to obtain the same number of clusters as for the entire transcriptome, and plotted the resulting clusters in an unsupervised manner onto a t-SNE plot calculated from the entire transcriptome (Fig. 1B-D). A qualitative inspection of the plots showed that the clusters resulting from a ‘random gene set’ did not clearly coincide with any clusters identified using the entire transcriptome, suggesting that they failed as transcriptional predictors for any given cell type (Fig. 1A, B). By contrast, ‘transcription factor’ clusters showed good correspondence to our whole transcriptome clustering (Fig. 1A, C). Intriguingly, we found that the ‘core matrisome’ was sufficient to identify several cell type clusters (Fig. 1A, D). For example, ‘core matrisome’ clusters m-7, m-11, m-14, and m-17 corresponded roughly to skeletal progenitors (t-15), joint progenitors (t-3), skin (t-1), and vessel (t-10) clusters, as identified by the entire transcriptome (Fig. 1A, D). Thus, these core matrisome-identified clusters largely reflected cell types of tissues that are embedded in collagen-rich ECMs.

**Figure 1.**
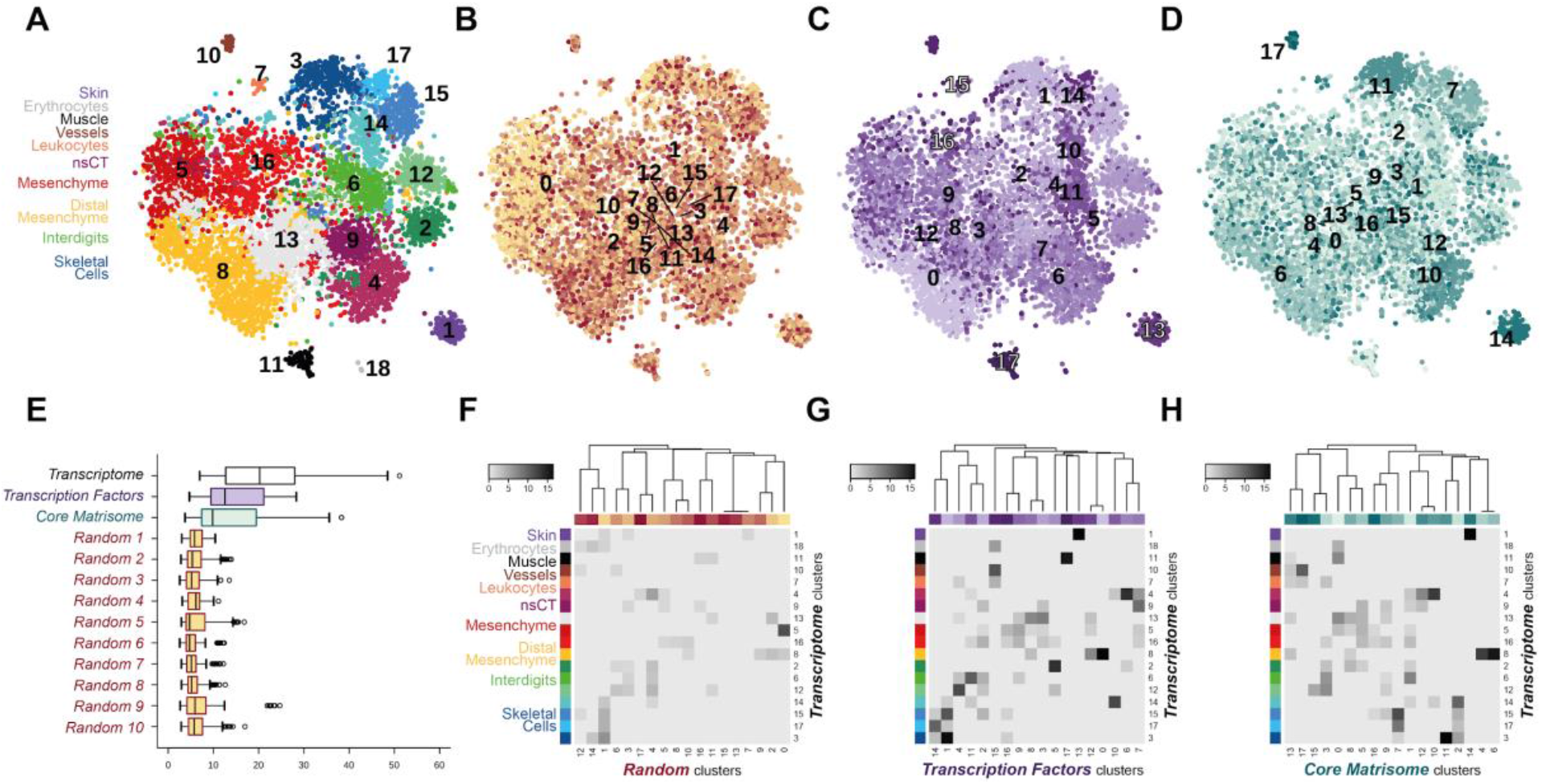
Core matrisome and transcription factors re-capitulate entire transcriptome cell clusters. (A) tSNE representation of 6823 HH29 chicken hindlimb autopod cells from Feregrino et al., 2019. Colors represent unsupervised clustering results based on the transcriptome (A), a randomly sampled set of genes (B), transcription factors (C), and the core matrisome (D). (E) Boxplot of Euclidean distances between clusters based on average expression of 2000 variably expressed genes, calculated for transcriptome, transcription factor, core matrisome, and 10 sets of random gene clusters. (F-H) Heatmap of square root of negative log 10 probability of cluster overlap by hypergeometric test between ‘transcriptome’ and ‘random clusters’ (F), ‘transcription’ factor’ (G), and ‘matrisome’ (H). ‘Transcriptome’ clusters are grouped by tissue or cell type. For (A-F) details, see Data Source File 1.

To quantify the separation among random genes-, transcription factors-, and core matrisome-based clusters, we plotted the distribution of all pairwise Euclidean distances, i.e. distances between all pairs of clusters, and compared them to the entire transcriptome result. Both the ‘core matrisome’ and the ‘transcription factors’ clusters clearly outperformed ten iterations of ‘randomly-picked’ genes subsets of equal size (Fig. 1E). Moreover, using a hypergeometric test, we were able to demonstrate that the probability of cluster overlap -between the entire transcriptome clusters and the three subsets clusters -was substantially higher for ‘core matrisome’ and the ‘transcription factors’ clusters (Fig. 1F-H). For the ‘core matrisome’, this was particularly evident for clusters corresponding to cell types known to produce a complex ECM, such as skeletal cells or skin (Fig. 1G). Moreover, even within the same cell type, the matrisome seemed able to distinguish discrete cell states. For example, ‘core matrisome’ clusters m-4 and m-6 reconstituted ‘transcriptome’ cluster t-8, the distal mesenchyme, indicating that the highly proliferative state of this mesenchymal sub-population is reflected by a distinct ‘matreotype’ (Fig. 1G). Taken together, our re-clustering analysis of chicken limb scRNA-seq data -using only the expression status of either core matrisome genes, transcription factors, or a random control gene set -indicates the potential of core matrisome gene expression status as a cell type and cell state identifier.

### Clustering performance and cell type identification by transcription factors and the core matrisome

To further assess the potential of such limited gene subsets to reliably identify cell types from scRNA-seq data, we next sought to quantify their ability to recreate our entire transcriptome cluster composition. We did this on a cluster-by-cluster as well as on a cell-by-cell basis. We first plotted -ordered by percentage -the respective cellular contributions of individual gene subset clusters to the 18 entire transcriptome clusters. As expected, ‘random genes’ clusters contributed almost uniformly to the different ‘transcriptome’ clusters (Fig. 2A). The median percentage contribution of the single-largest ‘random genes’ clusters -highlighted in yellow -was 17%, again reflective of that gene subset’s low information content regarding cell type identification. Certain ‘transcription factor’ clusters, however, contributed more than 90% of a given ‘entire transcriptome’ cluster (Fig. 2B). For example, ‘transcriptome’ cluster t-11, *i*.*e*., “muscle”, was represented to 99% by ‘transcription factor’ cluster tf-17. However, the same “muscle” cluster was only re-captured to 12% by the largest ‘random’ cluster contributor r-3 (compare Fig. 2A to B, ‘transcriptome’ cluster t-11). Likewise, the ‘matrisome’ gene subset also performed better than ‘random’, with the ‘Muscle’ cluster represented to 66% by ‘matrisome’ cluster m-0, or ‘Skin’ recaptured to 91% by cluster m-14 (Fig. 2C). However, when comparing the ‘transcription factor’ and ‘matrisome’ clustering performances within the closely lineage-related lateral plate mesoderm-derived cell types, differences between the two gene subsets emerged. Lateral plate mesoderm-derived tissues in our sample included non-skeletal connective tissue (cl. t-4, t-9), undifferentiated mesenchyme (cl. t-13, t-5, t-16, t-8), interdigital mesenchyme (cl. t-2, t-6, t-12) and skeletal progenitors (cl. t-14, t-15, t-17, t-3). Amongst these, certain cell type clusters contributing to mesenchymal tissues were well defined by their ‘transcription factor’ signature, yet much less so by their ‘matrisome’ expression status (e.g. compare cl. t-8, t-2, t-12, Fig. 2B and C). Again, some of these discrepancies might relate to the fact that ‘matrisome’ signatures can also be indicative of different cell states, whereas ‘transcription factors’ profiles assign predominantly to cell types. However, cell types contributing to more differentiated tissues with complex ECM composition were equally well defined by both ‘transcription factor’ and ‘matrisome’ gene expression signatures (*e*.*g*., cl. t-4, and t-14, t-15, t-17, t-3).

**Figure 2.**
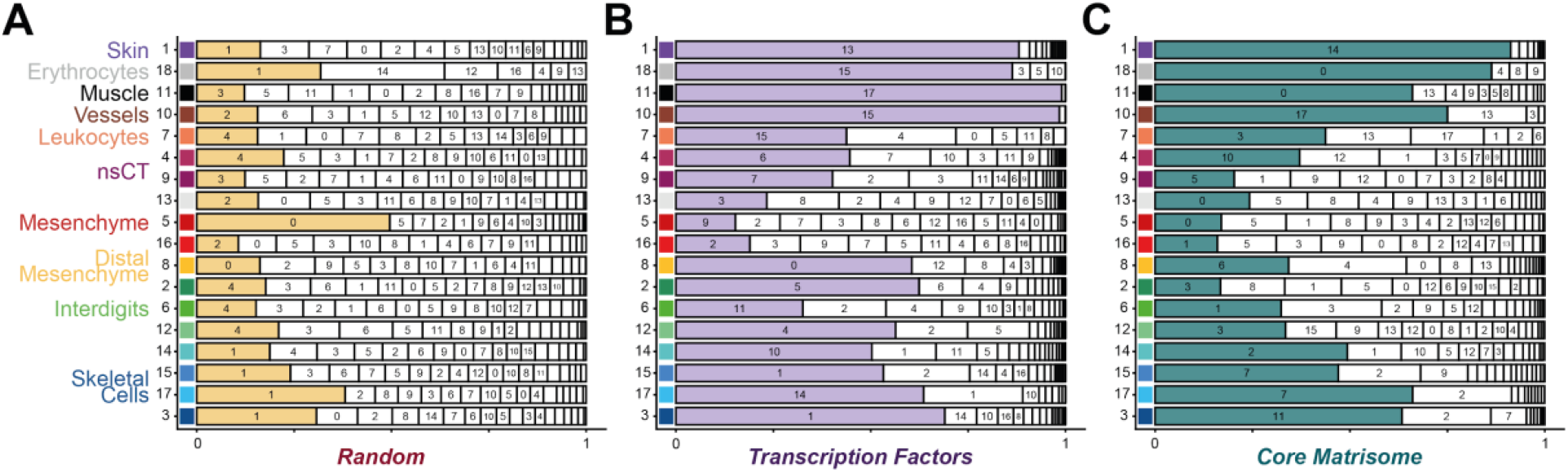
Relative cluster contributions to transcriptome clusters. (A) Relative contribution of ‘random’ clusters to the ‘transcriptome’ clusters ordered by size. Cluster IDs are indicated where possible and the biggest contribution per transcriptome cluster is highlighted in color. (B) ‘transcription factor’ cluster contributions and (C) ‘matrisome’ cluster contributions are indicated in the same manner. ‘transcriptome’ clusters are grouped by cell type.

### The core matrisome predicts preferentially ECM-rich cell types in early development

To quantify the ability of all our three gene-input-subsets -‘random’, ‘transcription factor’, and ‘matrisome’ -to correctly predict cluster membership of our “Gold Standard” transcriptome clustering, we decided to use a binary classification scheme based on pairs of cells being in the same or different clusters [29]. Each pair of cells was classified as either “true positive” (TP: two cells are in the same cluster regardless of the input data used), “true negative” (TN: two cells are in different clusters regardless of input data), “false positive” (FP: two cells are in the same cluster although they are in different clusters in the “Gold Standard”), and “false negative” (FN: two cells are in different clusters although they are in the same cluster in the “Gold Standard”) (Fig. 3A). Based on the cumulative numbers of TP, TN, FP, and FN of these binary cell pair classifications, we then calculated three different indices commonly used to compare different clustering algorithms [29]: the Rand index, also known as accuracy (*R* index), which measured the percentage of correct classifications; the Jaccard index of overlap (*J* index), which was calculated as the intersection of the two sets divided by the union of the two sets; and the Fowlkes-Mallows index (*FM* index), which represented the geometric mean of precision and recall. The closer each of these indices scores to 1, the more similar the respective gene subset clustering can be considered to the transcriptome “Gold Standard” clustering. Regardless of the index used, the ‘core matrisome’ and ‘transcription factor’ gene subsets clearly outperformed ten iterations of ‘random genes’, with ‘transcription factors’ scoring slightly higher than ‘matrisome’ genes (Fig. 3B).

**Figure 3:**
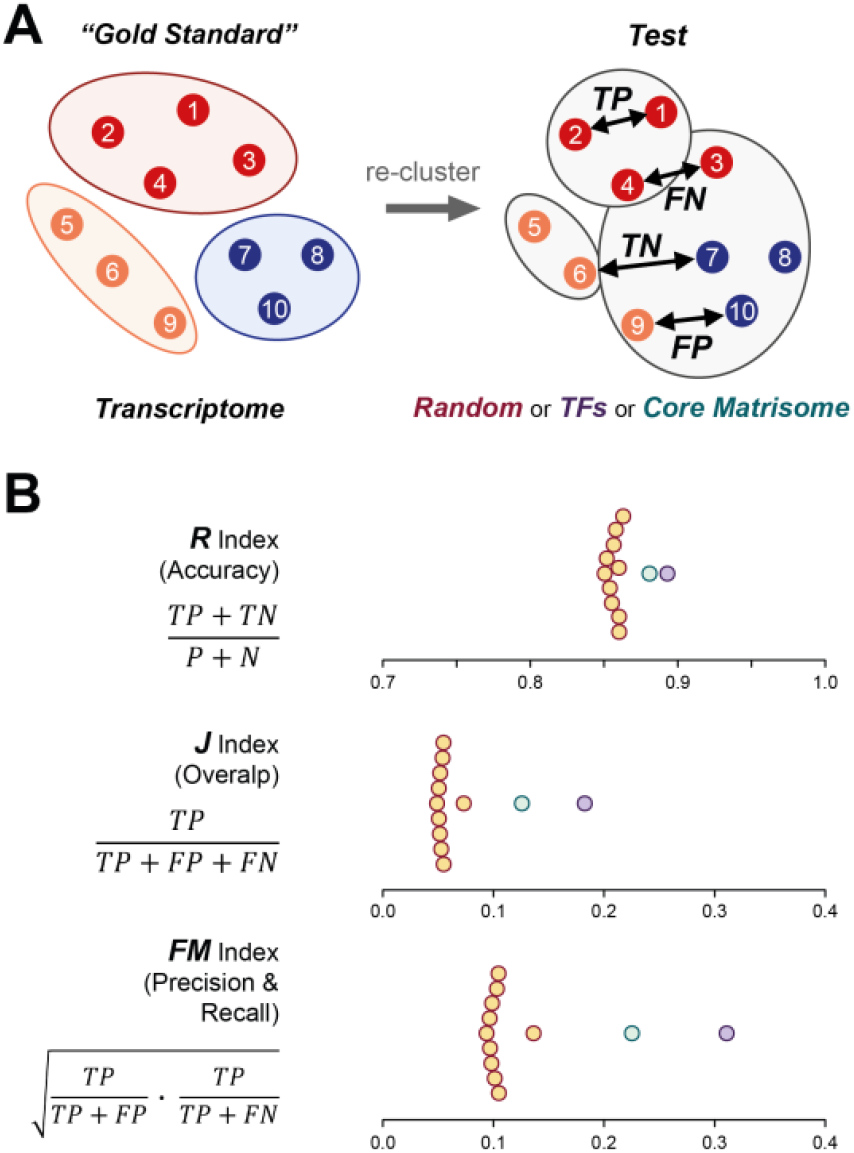
Binary classification and re-cluster indices. (A) Each pair of cells in a given ‘Test’ clustering -i.e. ‘Core Matrisome’, ‘Transcription Factor’ or ‘random’ -is classified based on their relationship to the “Gold Standard” clustering, as calculated from the entire transcriptome. (B) Based on those binary classifications the quality of each ‘Test’ clustering is measured with the three indices following Kafieh and Mehridehnavi, 2013. ‘Test’ clustering indices are calculated for transcription factors (purple), core matrisome (green), and 10 random gene set clusterings (yellow). TP: true positive, TN: true negative, FP: false positive, FN: false negative, R: Rand index, J: Jaccard index, FM: Fowlkes-Mallows index.

Collectively, we demonstrate that both ‘transcription factors’ and ‘matrisome’ genes can be used as cell type identifiers in scRNA-seq data. The extent to which this holds true, however, seems to depend on the tissue type to which the respective cell types contribute, differences in cell state, as well as the ontogenetic state of their differentiation.

### Predictive power of ‘core matrisome’ signature depends on developmental differentiation and is evolutionarily conserved amongst vertebrates

To determine the effect of developmental progression on the cell type-predictive powers of the matrisome, we next focused our attention on an embryonic scRNA-seq times series. We incorporated a previously published time series of the developing mouse hind limb by Kelly and colleagues into our analysis [30]. In their scRNA-seq data sets, we found 244 to 254 out of the total 274 mouse core-matrisome genes expressed. Initial clustering of single-cell transcriptomes showed -as expected -similar tissue composition as in our chicken hindlimb data, as well as as an increase in cell type complexity from the earliest stage E11.5 to E18.5 (Fig. 4A, Data Source File 1).

**Figure 4.**
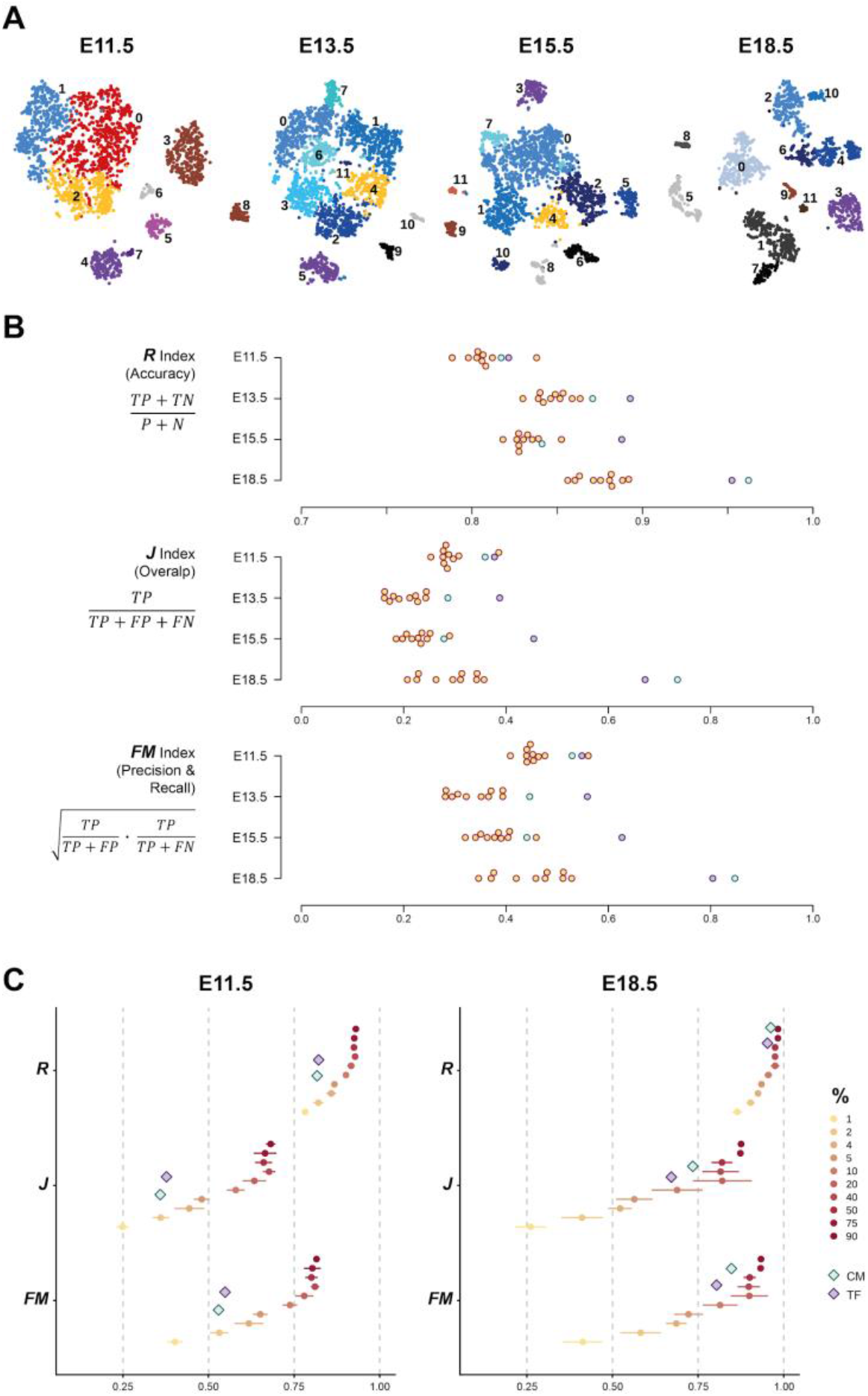
Predictive power of the matrisome during murine development. (A) tSNE representations of four mouse embryo limb data sets of increasing developmental stages (E11.5 to E18.5, E: embryonic day). Colors and numbers indicate ‘transcriptome’ clusters. Cluster annotation in Data source File 1. (B) Rand, Jaccard and Fowlkes-Mallows indices for all four stages. Colors as in Fig. 3B. (C) Indices of ‘matrisome’ and ‘transcription factor’ clustering (triangles) in E11.5 and 18.5 compared to indices of random gene sets of increasing size. Gradient indicates the percentage of the whole transcriptome represented by the random gene set. For each percentage, 5 gene sets were sampled independently and used for clustering.

Using the clusters identified by the entire transcriptome as a benchmark, we then re-clustered the data using either ‘random’, ‘transcription factor’ or ‘core matrisome’ gene sets of equal size, and compared their performances using the previously introduced indices. Over the course of the sampled developmental time window, the predictive powers of both ‘transcription factor’ and ‘core matrisome’ increased, *i*.*e*., more cell pairs were correctly attributed together in a way reflective of the entire transcriptome cell type clustering (Fig. 4B). To quantify this effect and relate it to the predictive powers of different fractions of ‘random’ genes from the entire transcriptome, we focused our attention on the two temporal extremes of the time series, E11.5 and E18.5. We randomly sampled increasing numbers of ‘random’ genes -from 1% to 90% of the entire transcriptome, each sampled and re-clustered 5 times -and plotted the spread of their performances in relation to the ‘transcription factor’ and ‘matrisome’ gene subsets (Fig. 4C, D). At E11.5, both ‘transcription factor’ and ‘matrisome’ gene subsets clusterings performed at about the rate of 2% of all genes, randomly selected from the transcriptome (Fig. 4C). At E18.5, however, their predictive powers had increased to a level of more than 10% of the entire transcriptome (Fig. 4D). This is noteworthy, as the number of expressed ‘cosre matrisome’ genes at that stage corresponds to only 1.28% of the entire transcriptome. Moreover, at these later stages, the ‘matrisome’ genes subset outperformed ‘transcription factors’ in these indices, likely a reflection of the ongoing maturation of tissues with high ECM content. We concluded, as tissues and their ECM mature, the expression status of the core matrisome becomes progressively better at delineating cell types.

## Discussion

Understanding the molecular parameters that define different cell types and states is fundamental to developmental and regenerative biology. Here we show that the expression status of a small subset of genes, the core matrisome, can suffice to identify cell types and states in the developing chick and mouse limb. Even though it corresponds to less than 2% of the entire transcriptome, we demonstrate that core matrisome expression encodes enough information to cluster scRNA-seq data according to cell types, and cell states. The predictive power of the matrisome increases with developmental time, and can even outperform transcription factors in more differentiated cell and tissue types with high ECM content.

These findings make sense with regards to developmental progression and tissue maturation. During ontogenetic development, transcription factors are thought to guide early differentiation trajectories and eventually specify terminally differentiated cell types [2,4,6]. At later stages of development, the ECM becomes increasingly important, instructing stem cell differentiation and regulating cell and tissue shape, morphogenetic movements, and organogenesis [16]. This holds especially true for tissues with complex ECM composition or high ECM turnover. Consistent with this, we found that in our chicken limb data, skin cells, muscle and skeletal progenitors clusters segregate especially well using core matrisome expression alone (Fig. 1 and 2). The lack of a clear ‘matrisome’-based clustering for some of the other mesenchymal cell populations may indicate a less specialized extracellular matrix, or, alternatively, the presence of different cell states within a cell type, each with its own putatively distinct ‘matreotype’. Moreover, the overall predictive power of the matrisome increases when comparing cell populations in ECM-rich tissues at progressively later stages of development (Fig. 4).

Previous work using scRNA-seq to determine molecular changes during adipogenesis (day 1-7) *in vitro* found that at day 3 the cell clustering was mainly driven by ECM genes, and at day 7 the core matrisome was one of the top ten most differentially expressed gene ontology terms [31]. Beyond development, planarian scRNA-seq revealed that muscles produce most of the matrisome, and inhibiting one key matrisome gene (hemicentin) resulted in severe epidermal ruffling and displacement of cells during homeostatic tissue turnover, suggesting an important role for tissue regeneration [26]. Furthermore, in healthy human lumbar discs, the core matrisome can be used to distinguish primary annulus fibrosus and nucleus pulposus cells based on 90 out of the 274 core matrisome genes being differentially expressed in the opposite direction using scRNA-seq [32]. Similarly, 115 matrisome genes are characteristically expressed in the six cell types that make up the human cutaneous neurofibroma microenvironment [33]. Beyond cell type distinction in tissues, a differential expression of matrisome genes can be observed when cells change their state from a healthy to a diseased cell. For instance, differential expression of matrisome genes was one main characteristic of reprogramming from normal fibroblasts, pericytes, and endothelial cells into tumor cells [34]. During cancer progression, deregulation of matrisome genes is a crucial step observed in early and late metastasis [35]. Moreover, core matrisome genes help identify circulating tumor cells in the blood using scRNA-seq [36,37]. These results support our conclusion that matrisome gene expression can serve as a key signature to determine individual cell types, as well as cell states.

This raises the question of why the matrisome is such a good predictor of cell type and state. It is well known that cells can be distinguished based on cell surface receptors [38]. However, it is less appreciated that each cell type can synthesize its own ECM that entails it with unique physical properties [15,39,40]. For instance, placing primary preadipocytes into decellularized ECMs derived from subcutaneous, visceral, or brown adipose tissue influences the preadipocytes’ terminal differentiation [41]. Hence, the physical properties of ECM seem to be able to dictate cellular fate and drive stem cell differentiation into neurons, muscle, or bone cells [42]. Besides providing instructive cues during development, ECMs can also change cellular status. Placing senescent cells or aged stem cells in a “younger ECM” rejuvenates these old cells to regain proliferative capacities or stem cell potential, respectively [43,44]. Similarly, placing tumor cells into an embryonic ECM reprograms to non-tumorigenic cells [45]. Hence, there is an intrinsic crosstalk between the ECM and the cells it encapsulates. ECMs, or niches, are made and adapted according to the respective cellular needs or states. Disrupting the crosstalk between cancer and cancer-associated fibroblasts, for instance, by a small molecule that inhibits chromatin remodeling and change matrisome gene expression (*i*.*e*., altering the matreotype), prevented tumor growth in xenograft mouse models [46]. Although we lack a current understanding of these underlying molecular crosstalk, these snapshots of ECM compositions -or matreotypes -clearly can reflect distinct cellular properties.

Accordingly, since matreotypes mirror cellular status, they also hold potentially promising prognostic value. For instance, 43 out of the 274 core matrisome genes are significantly upregulated across multiple cancer types, and 9 ECM genes predicted cancer outcome [47]. Another classifier similar to the matreotype concept is termed tumor matrisome index, which is based on 29 matrisome genes, reliably predicts low-and high-risk groups and chemotherapy responses for small cell lung cancer patients [48]. Matreotypes reflecting chronological age have been recently used to predict drugs that promote healthy aging [49]. Therefore, defining matreotypes has translational value for future biomedical research. Moreover, identifying different subpopulations of a given cell type will be critical to overcome the problem of cellular heterogeneity and aid personalized medical applications.

In summary, with our scRNA-seq analyses, we provided evidence for a previously postulated concept, namely that ‘each cell type produces its unique ECM’ [15,17]. While the best molecular proxies for cell-type identification continue to be discussed [1–3], we made the unexpected discovery that expressed core matrisome genes -corresponding to less than 2% of a typical transcriptome -hold enough information to re-cluster scRNA-seq data as well as transcription factor signatures. For more mature cells, the core matrisome embodied substantial predictive value to identify cell types and states. Hence, future work on defining matreotypes of different cell types and states might inform diagnostics and personalized medicine.

## Materials and Methods

### Matrisome gene lists

Curated matrisome gene lists for mouse and human are available on ‘The Matrisome Project’ (http://matrisome.org/; [14]. To create a matrisome list for chicken, a union of the human and mouse matrisome lists was used to define chick one-to-one orthologs in the ENSEMBL Galgal5.0 annotation.

### Single-cell RNA-sequencing data

Previously published single-cell RNA sequencing (scRNA-seq) datasets sampling the chicken embryonic limb [28] and the mouse embryonic limb [30]; stages E11.5 to E18.5) were used for all analyses. The raw data is accessible at Gene Expression Omnibus (GEO, https://www.ncbi.nlm.nih.gov/geo/), under accession numbers GSE130439 (chicken) and GSE142425 (mouse).

### Data pre-processing

Raw UMI count tables were used to initiate ‘Seurat objects’ for all mouse samples in R, using package Seurat v3.1.4. Next, low quality cells and outliers were filtered out. Chicken cells with an UMI count higher than 4 times the average UMI count, less than 20 percent of the median UMI count or more than 10 percent mitochondrial or ribosomal content were removed [28]. Mouse cells expressing less than 200, more than 6000 genes, or more than 10 percent mitochondrial RNA were removed. All expressed genes were considered. Normalization, identifying the top 2000 variable genes and scaling of the data was applied with Seurat’s built-in functions.

### Dimensionality reduction

For all chicken and mouse, Seurat objects, principal components analysis was performed on all expressed genes, and significant components were selected as such, if they were located outside of the Marchenko Pastur distribution [50]. The same criterion for significance was applied on all principal component analysis on core matrisome and random subset genes. The cells were visualized with the dimensionality reduction algorithm tSNE [51]. ‘matrisome’ and ‘random’ clusters for all datasets were represented on the same tSNEs generated from the ‘transcriptome’ principal components. To define a ‘gold standard’ of scRNAseq-based cell type clustering, k-nearest neighbour (kNN) graphs and Jaccard indices of overlap between a cell and its neighbours were used to create shared nearest neighbour (SNN) graphs, with the Seurat ‘FindNeighbors’ function using all expressed ‘transcriptome’ genes. Clusters of cells were then defined by ‘FindClusters’, by applying Louvain modularity optimization algorithms on SNN graphs. As the number of clusters can be influenced by the resolution parameters, please refer to the supplementary data for detailed parameters of significant dimensions and resolutions used in clustering for all samples. For ‘matrisome’-based clustering, all expressed core matrisome genes were considered for clustering. For ‘random’-based clustering, for ten iterations, randomly picked genes from the whole transcriptome were used such that they matched the number of expressed core matrisome genes, as well as resulting in the same number of individual clusters as the ‘transcriptome’ and ‘matrisome’ clustering. The core matrisome genes and transcription factors were excluded from this sampling. After ordering the transcription factors by expression variability, the set of top transcription factors matching the size of the expressed core matrisome was used to recluster the cells.

### Cluster cell type annotation

Differentially expressed genes between mouse clusters with a minimum natural log fold change of 0.25 were identified using a Wilcoxon rank sum test, and were then used to assign putative celltype identities of each cluster. Only genes expressed in at least 25% of cells in one of the two populations were considered. For all clusters, all and the top five differentially genes per cluster can be found in the Data Source File 1. Chicken clusters had been previously annotated (Feregrino et al., 2019).

### Distance Boxplots

To assess cluster-to-cluster proximity of ‘transcriptome’-, ‘matrisome’-, ‘random’-, and ‘transcription factor’-based clustering approaches, Euclidean pairwise distances between each cluster were calculated on the averaged scaled expression per cluster of the top 2000 variably expressed genes. The same 2000 genes were used to compare all three clustering approaches.

### Hypergeometric test

Probabilities of overlap between clusters were calculated with ‘phyper’. The hypergeometric test takes into account the size of the reference cluster (‘transcriptome’) *m*, the size of the test cluster (‘matrisome’, ‘random’, and ‘transcription factors’) *k*, the number of non-tested cells (total number of cells *N−m*) and the size of the overlap *x* to calculate the probability of the overlap to occur at random. Probabilities were calculated for overlaps between all clusters. Probabilities equal to zero were replaced with the smallest non-zero probability to prevent infinite values after transformation, and probabilities bigger than 0.05 were set to 1 for plot aesthetics. ‘Heatmap3’ [52] was used to plot square root negative log 10 transformed probabilities.

### Visualizing cluster contributions

Barplots were created with ‘ggplot2’ [53]

### Indices

The ‘Rand index’, also known as ‘accuracy’, was calculated as following:

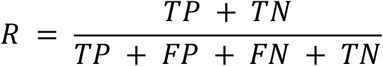

It measures the percentage of correct classifications.

The ‘Jaccard index of overlap’ is calculated as intersection over union. It does not take the TN into account, which represent the most classifications and might be confounding in the Rand index:

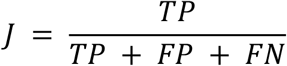

At last, the ‘Fowlkes-Mallows index’ is the geometric mean of precision and recall. Precision measures how many positive pairs (cells within the same cluster in the test clustering) are true positives (cells within the same cluster in the gold standard). Recall is the percentage of true positives identified by all actual positives:

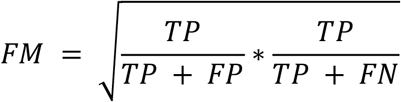

All indices range from 0 (no correct classification by the test clustering) to 1 (identical clustering by the test clustering).

### Comparing ‘matrisome’ and ‘transcription factors’ against ‘random’ subsets of increasing size

The information content of the ‘matrisome’ and the ‘transcription factor’ subset was compared to ‘random’ subsets containing 1, 2, 4, 5, 10, 20, 40, 50, 75, and 90 percent of the E11. 5 and E18.5 chicken transcriptomes. For each percentage, five ‘random’ subsets resulting in the same number of clusters as the ‘matrisome’ were sampled, clustered, and indices were calculated.

## Author contributions

FS, CF, PT and CYE designed the study. FS performed all analyses with help from CF. All authors interpreted the data. FS, PT, and CYE wrote the manuscript with comments from CF.

## Author Information

The authors have no competing interests to declare. Correspondence should be addressed to C. Y. E. and P.T.

## Acknowledgment

We thank Christian Beisel and the joint Genomics Facility Basel for help with single-cell sequencing, and members of both groups for useful discussions. All calculations were performed at sciCORE (http://scicore.unibas.ch/), the scientific computing center at the University of Basel. This work was supported by funding from the Swiss National Science Foundation PP00P3_163898 to CYE, and from the Swiss 3R Competence Centre (3RCC grant OC-2018-005), the Swiss National Science Foundation (SNSF project grant 310030_189242), and the University of Basel to PT.

