## Supplementary Table 1 for "Extracellular matrix gene expression signatures as cell type and cell state identifiers"

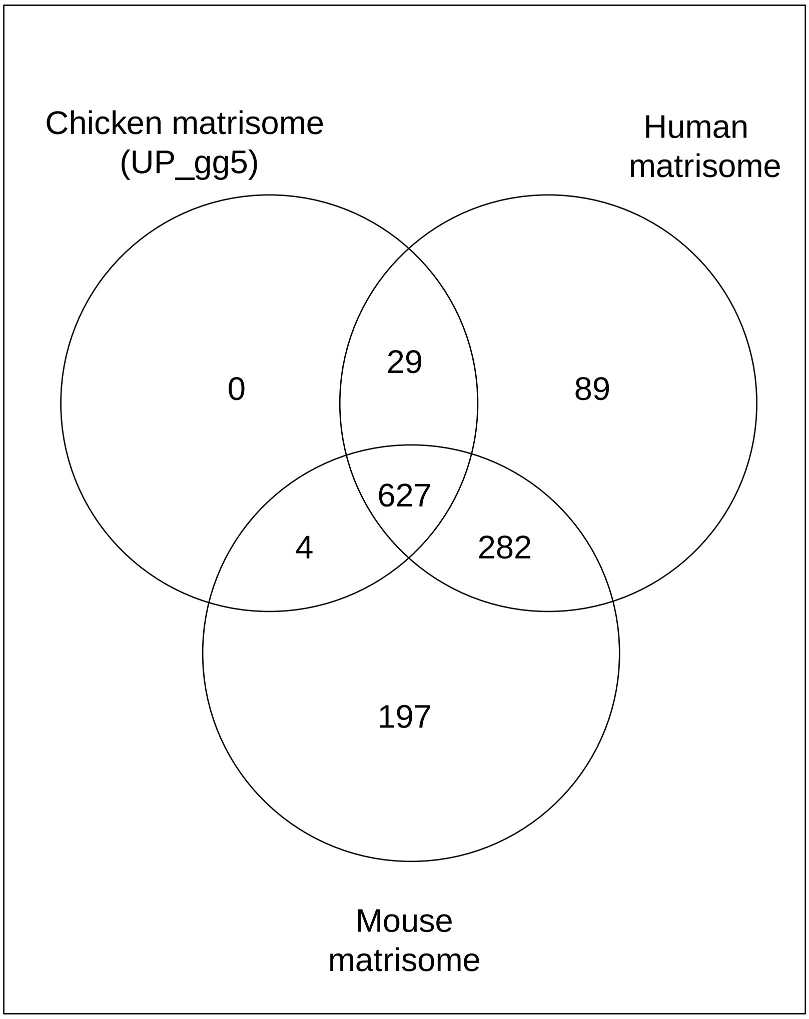


**Supplementary Figure 1: Overlap of our defined chick matrsiome with human and mice matrisome**

Venn diagram depicting overlap. For details see Supplementary Table 1 and Data Source File 1.
